# Genome-wide association study of habitual physical activity in over 277,000 UK Biobank participants identifies multiple variants including *CADM2* and *APOE*

**DOI:** 10.1101/179317

**Authors:** Yann C. Klimentidis, David A. Raichlen, Jennifer Bea, David O. Garcia, Lawrence J. Mandarino, Gene E. Alexander, Zhao Chen, Scott B. Going

## Abstract

**Background/Objectives:** Physical activity (PA) protects against a wide range of diseases. Engagement in habitual PA has been shown to be heritable, motivating the search for specific genetic variants that may ultimately inform efforts to promote PA and target the best type of PA for each individual.

**Subjects/Methods:** We used data from the UK Biobank to perform the largest genome-wide association study of PA to date, using three measures based on self-report (n=277,656) and two measures based on wrist-worn accelerometry data (n=67,808). We examined genetic correlations of PA with other traits and diseases, as well as tissue-specific gene expression patterns. With data from the Atherosclerosis Risk in Communities (ARIC; n=8,556) study, we performed a meta-analysis of our top hits for moderate-to-vigorous PA (MVPA).

**Results:** We identified 26 genome-wide loci across the five PA measures examined. Upon meta-analysis of the top hits for MVPA with results from the ARIC study, 8 of 10 remained significant at p<5×10^−8^. Interestingly, among these, the rs429358 variant in the *APOE* gene was the most strongly associated with MVPA. Variants in *CADM2*, a gene recently implicated in risk-taking behavior and other personality and cognitive traits, were found to be associated with regular engagement in strenuous sports or other exercises. We also identified thirteen loci consistently associated (p<0.005) with each of the five PA measures. We find genetic correlations of PA with educational attainment traits, chronotype, psychiatric traits, and obesity-related traits. Tissue enrichment analyses implicate the brain and pituitary gland as locations where PA-associated loci may exert their actions.

**Conclusions:** These results provide new insight into the genetic basis of habitual PA, and the genetic links connecting PA with other traits and diseases.

## Introduction

A physically active lifestyle has been shown to protect against a wide range of diseases, including cardiovascular disease, cancer, type-2 diabetes, osteoporosis, and Alzheimer’s disease ^1–4^. Levels of engagement in physical activity (PA) vary across individuals, and most people do not meet recommended levels to achieve health benefits. Although cultural, economic, and other environmental factors influence PA engagement ^5,6^, genetic factors also likely play a role. Understanding the genetic factors underlying inter-individual variation will better inform efforts to promote PA and potentially allow targeting the best type of PA for each person, what might be called “Precision Exercise Prescription”.

Evidence of genetic factors underlying the propensity to exercise in humans has been demonstrated in a number of studies ^7–14^. Several studies have utilized a candidate gene approach to identify specific genetic variants associated with a proclivity towards PA ^8,15–19^. This work generally focused on genes related to the serotonin and dopamine systems, energy metabolism, and neurotrophic factors. However, to our knowledge there have been only two previous reports of genome-wide association studies (GWAS) of PA ^20,21^, neither of which identified a locus at genome-wide significance, likely due to relatively small sample sizes. Thus, while previous work strongly suggests a genetic basis for engagement in PA, the genes that contribute to this healthy lifestyle behavior remain unknown.

In this study, we conduct the largest GWAS of PA to date, aiming to identify genetic variants associated with self-reported and accelerometry-based levels of habitual, leisure-time PA. We sought to identify variants in the UK Biobank, a large cohort study of 500,000 adults measured across a wide range of characteristics including genome-wide markers. We then examined the genetic correlation of PA with other traits, examined putative tissues where PA genes may exert their effects, and meta-analyzed the identified loci for MVPA with data on self-reported PA in an independent cohort from the Atherosclerosis Risk in Communities (ARIC) study.

## Methods

### Studies

Data from the UK Biobank study were used for discovery of variants. Briefly, the UK Biobank is a large prospective cohort study of approximately a half-million adults (ages 40-69) living in the United Kingdom (UK), recruited from 22 centers across the UK ^22^. All participants provided written informed consent. We also used data from the ARIC study, which is a prospective cohort study of over 15,000 adults aged 45-64 years that took place in four United States communities. Details of the ARIC study can be found elsewhere ^23^. To reduce the potential for confounding by population stratification, we included only individuals of white race/ethnicity in both studies.

### Physical activity

In the UK Biobank, self-reported levels of physical activity during work and leisure time were measured via a touchscreen questionnaire, in a fashion similar to the International Physical Activity Questionnaire ^24^. For moderate PA (MPA), participants were asked: “In a typical WEEK, on how many days did you do 10 minutes or more of moderate physical activities like carrying light loads, cycling at normal pace? (Do not include walking)”. For vigorous PA (VPA), participants were asked: “In a typical WEEK, how many days did you do 10 minutes or more of vigorous physical activity? (These are activities that make you sweat or breathe hard such as fast cycling, aerobics, heavy lifting)”. For each of these questions, those who indicated 1 or more such days were then asked “How many minutes did you usually spend doing moderate/vigorous activities on a typical DAY”. Participants were asked to include activities performed for work, leisure, travel and around the house. We excluded individuals who selected “prefer not to answer” or “do not know” on the above questions, those reporting not being able to walk, and individuals reporting more than 16 hours of either MPA or VPA per day. Those reporting >3hr/day of VPA or MPA were recoded to 3 hours, as recommended ^25^. Moderate-to-vigorous PA (MVPA) was calculated by taking the sum of total minutes/week of MPA multiplied by four and the total number of VPA minutes/week multiplied by eight, corresponding to their metabolic equivalents, as previously described ^24,26^.

Since heritability has previously been shown to be higher for intense/vigorous physical activity ^13^, we also considered VPA on its own. Because the distribution of minutes/week of VPA was highly skewed and zero-inflated, we chose to dichotomize minutes/week of VPA into those who reported 0 days of VPA, and those reporting 3 or more days of VPA and also reporting a typical duration of VPA that is 25 minutes or greater.

We used responses to the question “In the last 4 weeks did you spend any time doing the following?” and follow-up questions assessing the frequency and typical duration of “strenuous sports” and of “other exercises”. The possible responses to the initial question were: ‘walking for pleasure’, ‘other exercises’, ‘strenuous sports’, ‘light DIY’, ‘heavy DIY’, ‘none of the above’, and ‘prefer not to answer’. We identified individuals spending 2-3 days/week or more doing strenuous sports or other exercises (SSOE), for a duration of 15-30 minutes or greater. Controls were those individuals who did not indicate spending any time in the last 4 weeks doing either strenuous sports or other exercises.

Also, in the UK Biobank, approximately 100,000 participants wore an Axivity AX3 wrist-worn accelerometer, as previously described ^27^. We examined two measures derived from up to seven days of accelerometer wear: overall acceleration average, and fraction of accelerations > 425 milli-gravities (mg)^27^. Since the variable that is available in the UK Biobank is the fraction < 425 mg, we subtracted 1 from this variable. The 425 mg cutoff was chosen because this corresponds to an equivalent of vigorous physical activity, as previously reported ^28^. For both accelerometry variables, we excluded individuals with less than three days (72 hours) of data, those not having data in each one-hour period of the 24-hour cycle, and outliers with values more than 4 standard deviations above the mean.

In ARIC, self-reported PA was assessed for sports/exercise, within the previous year, based on a modification of the Baecke questionnaire ^29,30^. The sport/exercise index is based on responses to 4 items: frequency of participation in sports/exercise; frequency of sweating during sports/exercise; a subjective rating of the frequency of participation in sports/exercise compared to others in the same age group; the sum of frequency, duration, and intensity of up to 4 sports/exercises.

### Genotypes

The majority of UK Biobank participants were genotyped with the Affymetrix UK Biobank Axiom Array (Santa Clara, CA, USA), while 10% of participants were genotyped with the Affymetrix UK BiLEVE Axiom Array. Detailed quality control and imputation procedures are described elsewhere^31^. Imputation was performed using a combined panel of the Haplotype Reference Consortium ^32^ and the UK10K haplotype resource ^33^. Since corrections for potential problems with the position assignment of the SNPs from the UK10K haplotype resource were not available at the time of analysis, we only included SNPs imputed from the Haplotype Reference Consortium. To minimize the possibility of confounding due to population stratification, only participants who self-identified as White/British (European-descent), and who were not population outliers based on principal components analysis (PCA) were included. Individuals were excluded based on unusually high heterozygosity or >5% missing rate, a mismatch between self-reported and genetically-inferred sex, and on relatedness (i.e. samples not used in primary PCA). SNP exclusions were made based on Hardy-Weinberg equilibrium (p<1×10^−6^), high missingness (>1.5%), low minor allele frequency (<0.1%), and low imputation quality (info<0.4). A total of approximately 11.7 million SNPs were used in analyses.

In ARIC, participants were genotyped with the Affymetrix Genome-Wide Human SNP Array 6.0 (Affymetrix, Santa Clara, CA, USA). Standard quality control procedures were implemented prior to imputation with IMPUTE2 ^34^, using all individuals in the 1,000 Genomes phase 1 integrated v3 release. Quality-control procedures consisted of excluding SNPs with minor allele frequency < 1%, with missingness > 10%, and SNPs out of Hardy-Weinberg equilibrium (p<1 × 10^−6^), and excluding individuals with SNP missingness > 10%. We used principal components for the European-ancestry group as provided by ARIC in dbGaP. Briefly, LD pruning resulted in 71,702 SNPs that were used to derive principal components. A total sample size of 8,556 participants was used in the analysis.

### Statistical analyses

For the continuous variables in the UK Biobank (MVPA and accelerometry variables) we created an adjusted phenotype corresponding to the residual of the regression of the following independent variables on the respective dependent PA variable: age, sex, genotyping chip, first ten genomic principal components, center, season (month) at center visit or wearing accelerometer (coded 0 for Winter, 1 for Fall or Spring, and 2 for Summer). In sensitivity analyses, we considered the inclusion of the additional following covariates: levels of physical activity at work (coded as 0 by default, 1 for ‘sometimes’, 2 for ‘usually’, and 3 for ‘always’), extent of walking or standing at work (coded similarly as previous variable), and the Townsend Deprivation Index (TDI; a composite measure of deprivation as previously described ^35,36^). We also considered a third model in which body mass index (BMI) was included as an additional covariate. Since the MVPA and fraction of accelerations > 425 mg variables exhibited skewed distributions, we inverse-normalized these variables prior to inclusion in the models. Model residuals conformed to the assumptions of normality and homoscedasticity. For the categorical variables, we used logistic regression, including the same set of covariates listed above. We also sought to identify variants consistently associated with PA across all five measures. We thus searched for variants associated in the same direction, with p<0.001 with each of the five PA phenotypes. The prior probability under the null hypothesis for a SNP association at this level with all four phenotypes and with the same direction of association is 0.005 × 0.0025^4^ = 1.95 × 10^−13^. However, it is important to note that these five phenotypes are correlated with each other.

Given the association that we identified with the rs428358 variant in *APOE* (see results below), we performed two additional analyses. First, we examined the associations with the *APOE* ε4 genotype, using this SNP along with the rs7412 SNP. Individuals with homozygous CC genotypes at both of these SNPs were classified as homozygous for the APOE ε4 allele. Individuals with homozygous CC genotypes at either SNP and heterozygous at the other SNP were classified as being heterozygous for the ε4 allele. We excluded individuals heterozygous at both SNPs. We assumed an additive model in association testing. Second, to examine whether this association may be driven by individuals with a known family history of Alzheimer’s disease increasing their levels of PA, we examined the association of a binary variable indicating any self-reported first-degree family history (mother, father, or siblings) of Alzheimer’s disease or dementia with MVPA.

All genome-wide significant loci for MVPA were examined in ARIC, where we modeled PA as a continuous variable (as described above). We used multiple linear regression to model PA as a function of: age, sex, first ten genomic principal components, BMI, center, season (coded in the same way as described above). Residuals from this model conformed to the assumptions of normality and homoscedasticity. They were standardized to have a mean of 0 and standard deviation of 1, and were used as the outcome in the genome-wide SNP association analysis. We performed meta-analysis of the top hits for MVPA in the UK Biobank with the corresponding SNP association results in ARIC, using fixed-effects inverse-variance weighted meta-analysis. All GWAS analyses were performed with PLINK 2.0, assuming an additive genetic model ^37^. Additional analyses were performed with R statistical software ^38^. To examine association of genetic variants with gene expression patterns in different tissues, we used the web-based platform, Functional Mapping and Annotation of Genome-Wide Association Studies (FUMA GWAS) ^39^, which uses data from GTEx ^40^.

We used the summary statistics from our GWAS to examine genetic correlation of PA with over 200 traits and diseases using LD score regression ^41,42^, implemented in an online interface (http://ldsc.broadinstitute.org/). The genetics of these other traits and diseases are inferred from previously published GWAS. A significant genetic correlation was considered to have a p< 2.5 × 10^−4^, assuming a correction for 200 different tests, which is conservative given that many of the traits/diseases tested are correlated with each other. LD score regression intercepts and chip heritability estimates were also recorded.

## Results

### Self-reported PA in UK Biobank

A summary of self-report PA variables can be found in Table 1. BMI and TDI were consistently negatively associated with these variables, whereas warmer season and male gender were consistently positively associated with them (see Supplementary Table 1). Physical activity at work was positively associated with MVPA and VPA, and negatively associated with SSOE. Self-report PA measures were weakly correlated with accelerometry-based measures (see Supplementary Table 2). ‘Chip heritability’ estimates for self-report PA measures were approximately 5-6% (Supplementary Table 3). LD score regression intercepts (<1.02) suggest no significant systematic inflation of test statistics. In over 277,000 individuals, we found ten loci significantly associated (p < 5 × 10^−8^) with MVPA and four loci with VPA (see Figure 1 and Table 2). Most notably, the C allele at SNP rs429358 in *APOE* is associated with higher self-reported MVPA. This allele is also nominally associated with higher levels of other PA measures (VPA: p=8.4 × 10^−6^; SSOE: p=0.049; average acceleration: p=3.7 × 10^−3^). Testing the association of the Alzheimer’s disease-related *APOE* ε4 allele with MVPA resulted in nearly identical findings. In models adjusted for other covariates, this *APOE* variant remained genome-wide significant (see Supplementary Tables 4 and 5). There were 33,337 individuals reporting any family history of Alzheimer’s disease or dementia among parents and siblings. These individuals reported lower levels of MVPA (p=1.2 × 10^−4^). An intronic variant in *CADM2* was associated with SSOE (see Table 2). A variant at the same locus was also associated with VPA in the model including all covariates (see Suppl. Table 5). Variants in and near *MMS22L* were identified with VPA and SSOE, respectively. Upon meta-analysis of the 10 top hits for MVPA with the results in ARIC, 8 were genome-wide significant (p<5 × 10^−8^), including the APOE variant (see Table 3). In models adjusting for additional covariates (PA at work, TDI, BMI), the results were similar, but with fewer loci reaching genome-wide significance (see Supplementary Tables 4 and 5 and Supplementary Figures 1 and 2). Notably, the *APOE* and *CADM2* loci remained significant after adjustment for other covariates.

**Table 1:**
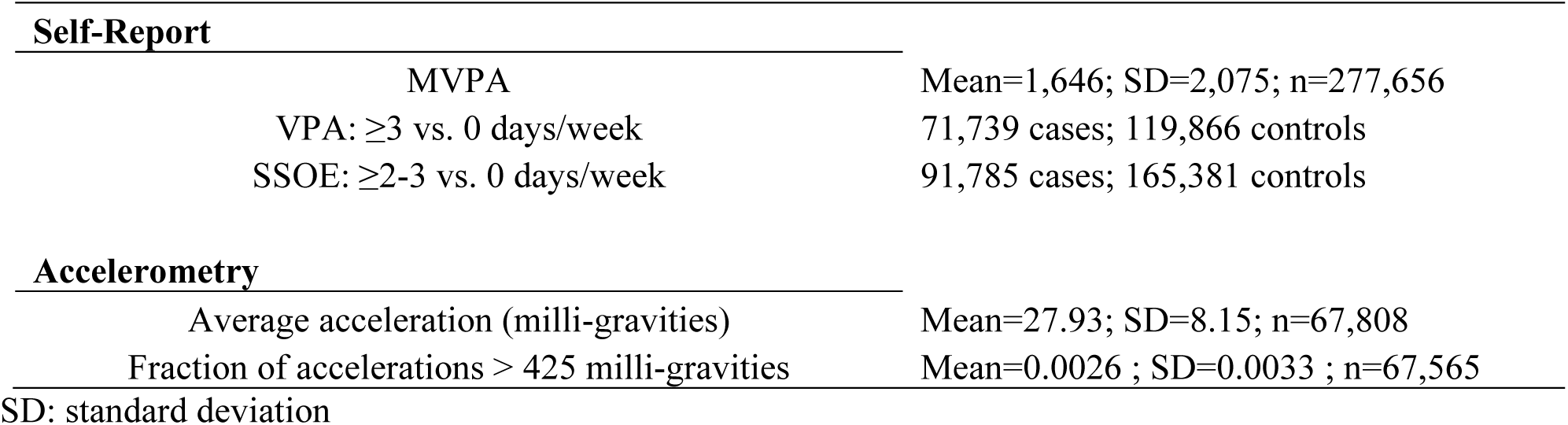
Summary of PA phenotypes in the UK Biobank.

**Table 2:** Summary of polymorphisms identified in the UK Biobank.

| SNP ID | Chr. | Gene/Nearest Gene | Position | EA | EAF | Beta/OR | p-value | n |
| --- | --- | --- | --- | --- | --- | --- | --- | --- |
| *MVPA* |  |  |  |  |  |  |  |  |
| rs429358 | 19 | *APOE* | 45,411,941 | T | 0.840 | -0.024 | 4.98E-11 | 277,656 |
| rs72737787 | 9 | *ZCCHC7* | 37,336,741 | G | 0.650 | -0.018 | 3.66E-10 | 277,656 |
| rs1043595 | 7 | *CALU* | 128,410,012 | G | 0.720 | 0.018 | 1.48E-09 | 277,656 |
| rs1921981 | 21 | *BACE2* | 42,422,547 | G | 0.670 | 0.017 | 4.03E-09 | 277,656 |
| rs3129981 | 6 | *HCG20* | 30,758,857 | C | 0.830 | 0.021 | 5.13E-09 | 277,656 |
| rs921917 | 7 | *C7orf72* | 50,228,738 | C | 0.510 | -0.016 | 1.19E-08 | 277,656 |
| rs12147808 | 14 | *C14orf64* | 98,618,705 | C | 0.790 | -0.019 | 1.30E-08 | 277,656 |
| rs124672 | 8 | *KCNK9* | 140,722,178 | A | 0.710 | -0.017 | 2.22E-08 | 277,656 |
| rs148854222 | 5 | *SLC1A3* | 36,505,944 | T | 0.997 | -0.143 | 2.87E-08 | 277,656 |
| rs12142550 | 1 | *HAX1* | 154,258,549 | C | 0.750 | 0.017 | 4.60E-08 | 277,656 |
| *Vigorous PA: ≥ 3 vs. 0 days/week* |  |  |  |  |  |  |  |  |
| rs6955240 | 7 | *EXOC4* | 133,581,873 | G | 0.610 | 1.044 | 9.57E-10 | 191,473 |
| rs3781411 | 10 | *CTBP2* | 126,715,436 | C | 0.880 | 1.063 | 4.88E-09 | 191,473 |
| rs6667222 | 1 | *HAX1* | 154,253,661 | A | 0.750 | 1.046 | 9.55E-09 | 191,473 |
| rs6909774 | 6 | *MMS22L* | 97,687,471 | A | 0.250 | 1.044 | 3.34E-08 | 191,473 |
| *Strenuous sports or other exercises: ≥ 2-3 vs. 0 days/week* |  |  |  |  |  |  |  |  |
| rs1376935 | 3 | *CADM2* | 85,236,425 | G | 0.690 | 0.955 | 1.28E-12 | 256,850 |
| rs10946808 | 6 | *HIST1H1D* | 26,233,387 | A | 0.730 | 0.962 | 3.10E-09 | 256,850 |
| rs1638525 | 17 | *AKAP10* | 19,848,594 | G | 0.600 | 1.035 | 7.90E-09 | 256,850 |
| rs35622985 | 6 | *MMS22L* | 97,783,799 | G | 0.680 | 0.964 | 8.81E-09 | 256,850 |
| rs705692 | 1 | *CAMTA1* | 7,480,217 | C | 0.110 | 0.947 | 1.53E-08 | 256,850 |
| rs1959759 | 14 | *DCAF5* | 69,632,877 | A | 0.360 | 1.035 | 1.62E-08 | 256,850 |
| rs4411372 | 13 | *STK24* | 99,130,423 | T | 0.720 | 0.964 | 2.36E-08 | 256,850 |
| rs181053839 | 2 | *WDPCP* | 63,486,141 | G | 0.996 | 0.767 | 2.51E-08 | 256,850 |
| rs9949626 | 18 | *LOC100131655* | 74,509,958 | C | 0.470 | 1.033 | 3.02E-08 | 256,850 |
| *Accelerometry – Average acceleration* |  |  |  |  |  |  |  |  |
| rs62055545 | 17 | *MAPT-AS1* | 43,964,561 | C | 0.780 | -0.038 | 1.41E-09 | 67,808 |
| rs34517439 | 1 | *FUBP1* | 78,450,517 | C | 0.880 | 0.044 | 4.23E-08 | 67,808 |
| rs4747438 | 10 | *DNAJC1* | 22,124,263 | C | 0.320 | -0.030 | 4.62E-08 | 67,808 |
| *Accelerometry – Fraction accelerations > 425 milli-gravities* |  |  |  |  |  |  |  |  |
| rs10851869 | 15 | *PML* | 74,331,083 | T | 0.570 | 0.027 | 3.66E-08 | 67,565 |

**Table 3:**
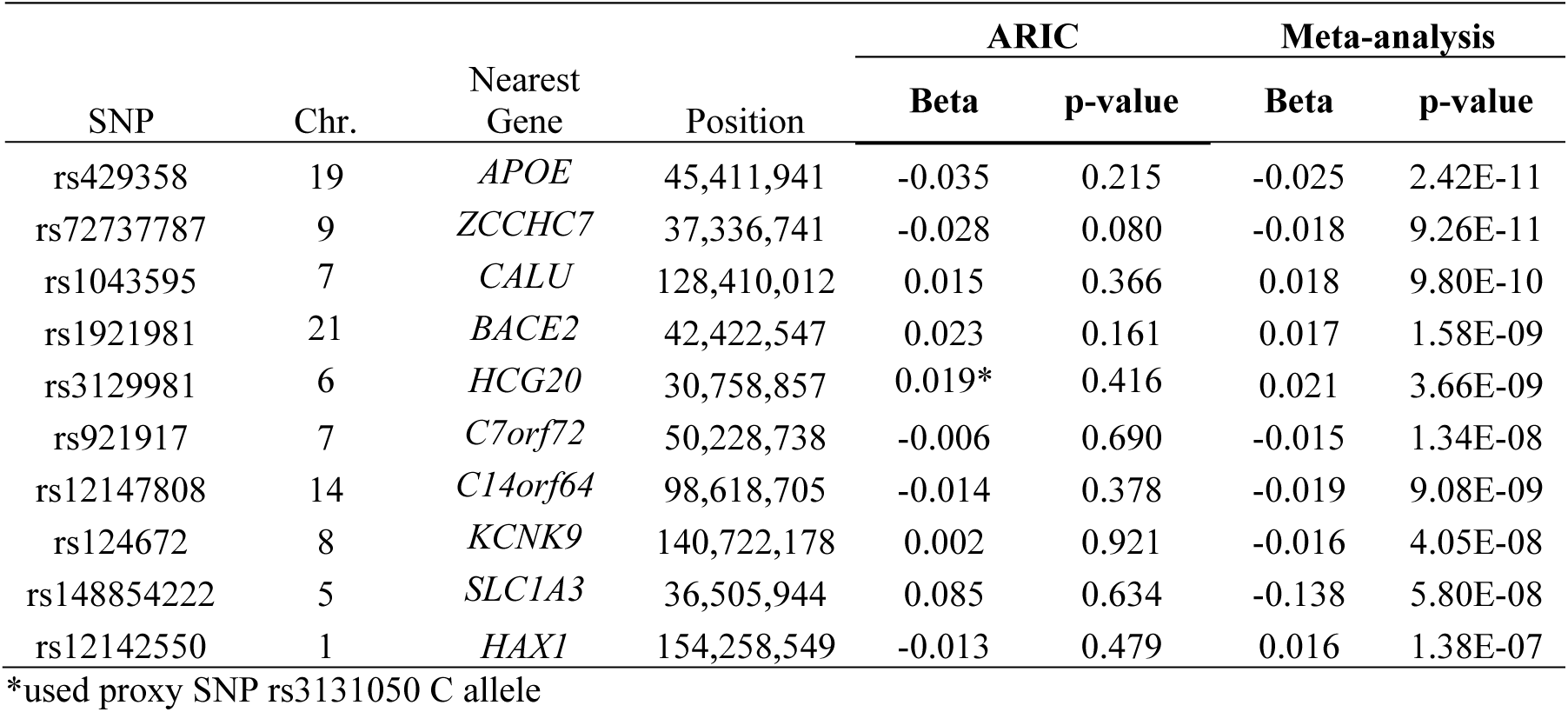
Meta-analysis of UK Biobank MVPA top hits with ARIC PA.

**Figure 1:**
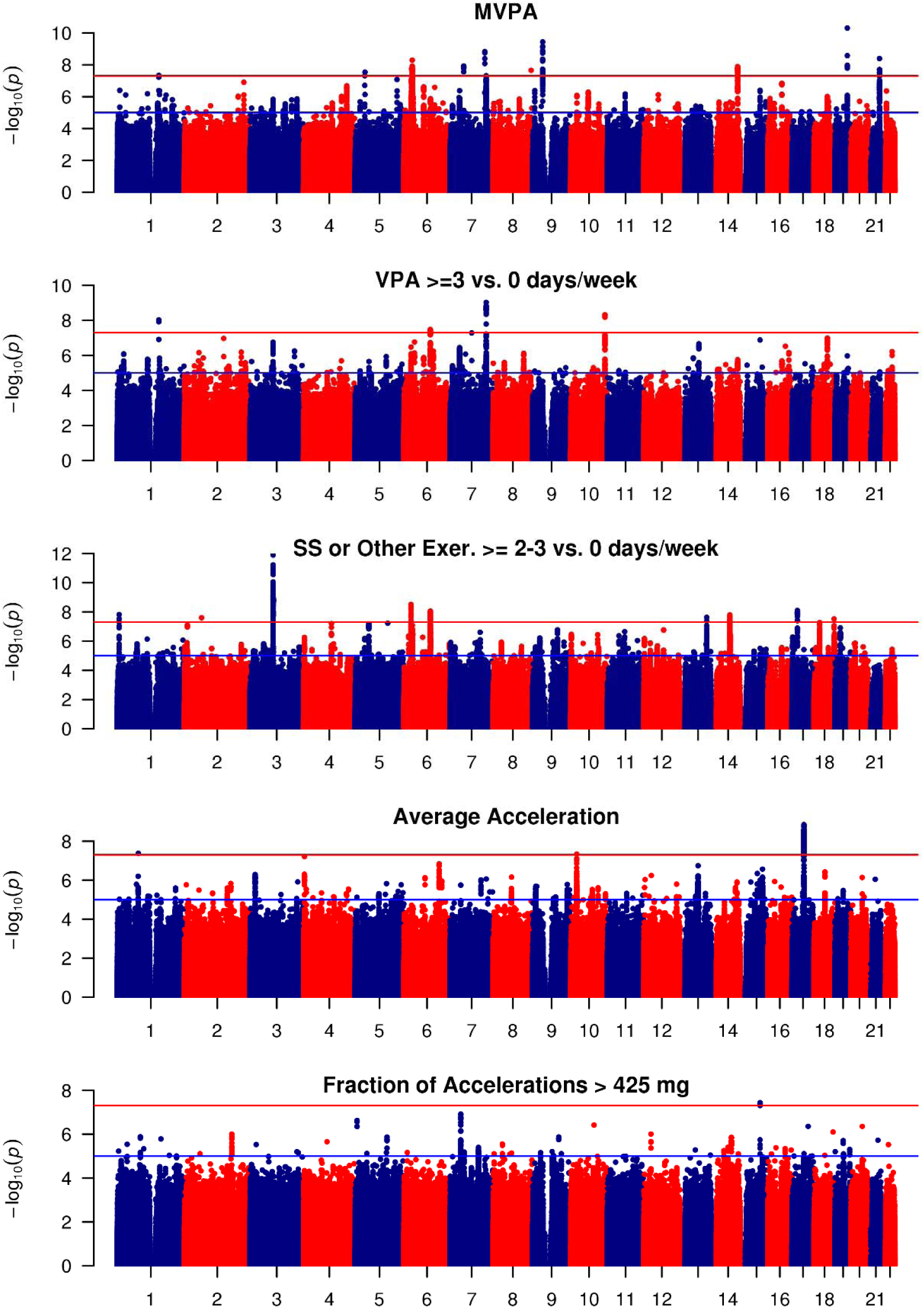
Manhattan plot of GWAS for self-reported MVPA and VPA, strenuous sports or other exercises (abbreviated as SS or Other Exer.), and for accelerometer-based average accelerations and fraction of accelerations > 425 mg. Negative log10-transformed p-value for each SNP is plotted by chromosome and position (x-axis). The red and blue horizontal lines represent thresholds for genome-wide significant and suggestive associations, respectively.

**Figure 2:**
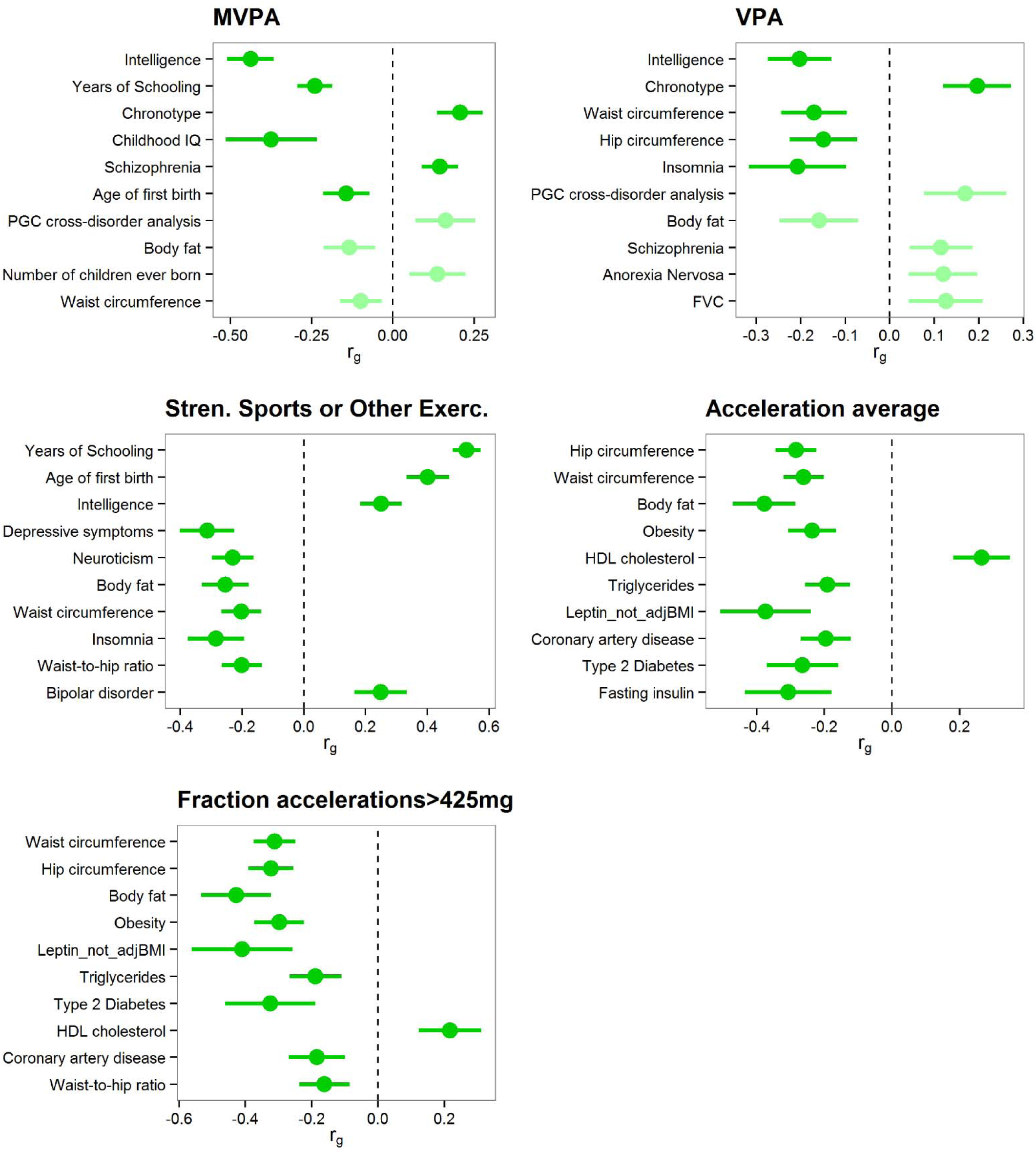
Genetic correlation of PA variables with other traits and diseases. Traits/diseases are ordered from top to bottom in order of increasing p-value for the ten traits/diseases with strongest degree (according to p-value) of genetic correlation with respective measure of PA. Horizontal position of bars corresponds to the genetic correlation (r_g_) between PA and the respective trait/disease. Error bars represent 95% confidence intervals for r_g_ estimates. Bright green bars represent traits that showed a correlation with p-value <2.5E-4, and light green bars represent traits with genetic correlation p<0.05. We excluded highly redundant traits (e.g. obesity, overweight) after leaving higher ranked one in.

### Accelerometer-based PA in UK Biobank

‘Chip heritability’ estimates for the accelerometry-based measures were higher (15% for average acceleration, and 11% for fraction of accelerations >425 mg) than for self-report PA measures (Supplementary Table 3). LD score regression intercepts (<1.009) suggest no significant systematic inflation of test statistics. Three loci were found to be significantly associated with average acceleration and one locus with fraction of accelerations >425 mg (see Table 2 and Figure 1). In models adjusted for other covariates, only the *MAPT_AS1* locus remained consistently genome-wide significant (see Supplementary Tables 4 and 5 and Supplementary Figures 1 and 2).

We found thirteen loci associated with each of the five PA measures at p<0.005, and with the same direction of association (Table 4). These include an intronic variant in *DNAJC1*, a variant just upstream of *DCAF5*, and a missense variant in *PML*.

**Table 4:** Loci consistently associated with PA across all five measures (each p<0.005).

| SNP | Chr. | Nearest Gene | Position | Effect Allele | MVPA |  | VPA |  | SSOE |  | AA |  | AF>425 |  |
| --- | --- | --- | --- | --- | --- | --- | --- | --- | --- | --- | --- | --- | --- | --- |
|  |  |  |  |  | Beta | p | OR | p | OR | p | Beta | p | Beta | p |
| rs4361077 | 2 | *ACVR1* | 158,556,835 | G | 0.009 | 3.70E-03 | 1.032 | 1.11E-04 | 1.022 | 2.25E-03 | 0.028 | 4.93E-06 | 0.018 | 1.76E-03 |
| rs10930438 | 2 | *LOC101926913* | 171,606,871 | A | -0.010 | 4.75E-04 | 0.973 | 2.38E-04 | 0.976 | 1.07E-04 | -0.022 | 9.09E-05 | -0.015 | 4.68E-03 |
| rs2113077 | 5 | *ISL1* | 50,799,442 | A | 0.009 | 1.07E-03 | 1.024 | 7.05E-04 | 1.029 | 2.32E-06 | 0.016 | 2.31E-03 | 0.016 | 1.08E-03 |
| rs1993246 | 7 | *KCCAT333* | 17,477,177 | C | -0.008 | 3.58E-03 | 0.976 | 5.39E-04 | 0.979 | 4.76E-04 | -0.016 | 2.74E-03 | -0.014 | 4.68E-03 |
| rs13284832 | 9 | *MAPKAP1* | 128,435,126 | A | -0.007 | 4.11E-03 | 0.975 | 1.76E-04 | 0.977 | 9.56E-05 | -0.021 | 4.06E-05 | -0.014 | 4.76E-03 |
| rs7910002 | 10 | *DNAJC1* | 22,050,570 | G | -0.009 | 8.73E-04 | 0.973 | 2.83E-04 | 0.982 | 4.47E-03 | -0.030 | 7.90E-08 | -0.020 | 1.03E-04 |
| rs9579775 | 13 | *ZMYM2* | 20,616,557 | A | 0.016 | 3.23E-05 | 1.045 | 1.82E-05 | 1.032 | 5.13E-04 | 0.023 | 2.95E-03 | 0.024 | 8.64E-04 |
| rs10135643 | 14 | *DCAF5* | 69,517,406 | A | 0.009 | 6.46E-04 | 1.027 | 1.72E-04 | 1.033 | 7.93E-08 | 0.019 | 4.57E-04 | 0.018 | 2.24E-04 |
| rs10145335 | 14 | *C14orf177* | 98,547,748 | G | -0.016 | 6.81E-08 | 0.973 | 3.98E-04 | 0.980 | 2.39E-03 | -0.018 | 3.03E-03 | -0.017 | 1.60E-03 |
| rs5742915 | 15 | *PML* | 74,336,633 | T | -0.012 | 3.67E-06 | 0.965 | 1.33E-07 | 0.973 | 3.14E-06 | -0.022 | 2.44E-05 | -0.022 | 3.04E-06 |
| rs185231044 | 17 | *RHBDL3* | 30,637,986 | G | -0.038 | 2.42E-04 | 0.921 | 2.38E-03 | 0.920 | 3.82E-04 | -0.064 | 1.97E-03 | -0.060 | 1.73E-03 |
| rs113351744 | 18 | *LINC01029* | 75,585,414 | G | -0.030 | 4.59E-03 | 0.908 | 3.58E-04 | 0.899 | 5.77E-06 | -0.063 | 2.30E-03 | -0.058 | 2.43E-03 |
| rs12460611 | 19 | *CCNE1* | 30,326,600 | A | 0.009 | 2.15E-03 | 1.029 | 1.25E-04 | 1.026 | 4.69E-05 | 0.026 | 4.01E-06 | 0.021 | 2.94E-05 |

### Genetic correlation and tissue enrichment analysis

We examined the genetic correlation between habitual PA and over 200 other traits/diseases. We found highly significant negative genetic correlations of MVPA and VPA with intelligence (see Figure 2). We also found significant positive genetic correlations with early-morning chronotype and schizophrenia. Interestingly, however, we found a positive correlation of SSOE with years of schooling and intelligence, as well as with age at first birth and bipolar disorder. SSOE was negatively genetically correlated with depressive symptoms, neuroticism, body fat, waist circumference, and insomnia (see Figure 2). Among the accelerometry-based variables, we found highly significant negative genetic correlations of PA with waist and hip circumference, body fat, obesity, BMI, and triglycerides (see Figure 2). Genetic correlation results remained very similar with GWAS models including activity at work and TDI as covariates (see Supplementary Figure 3), except for generally attenuated correlations with intelligence in the model with all covariates except BMI. Upon addition of BMI as a covariate, the direction of genetic correlation between PA and obesity traits was reversed (see Supplementary Figure 4). As we note below, caution may be warranted in interpreting results from these adjusted models. In ARIC, we found similar genetic correlations of PA with waist circumference (r_g_=−0.13), chronotype (r_g_=0.04), years of schooling (r_g_=0.16), and schizophrenia (r_g_=0.32), although these do not reach statistical significance.

**Figure 3:**
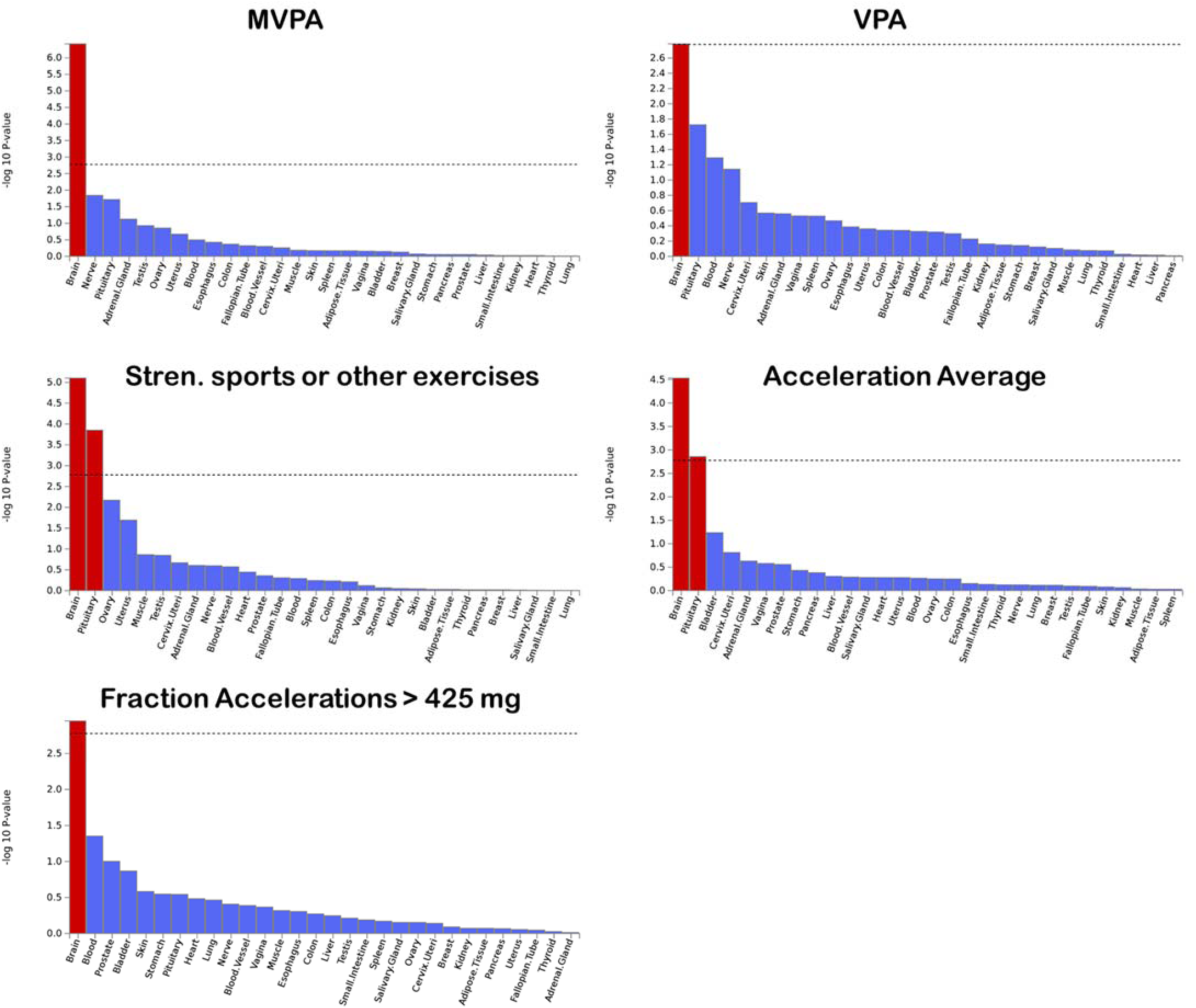
Results of tissue enrichment analysis using eQTL results from GTEx RNA-seq data for PA-associated loci. Dashed line represents the Bonferroni-corrected significance threshold.

Tissue enrichment analysis using eQTL data from GTEx implicate the brain and pituitary gland as primary tissues through which the PA-associated loci may exert their effects (see Figure 3 and Supplementary Figures 5 and 6).

## Discussion

Given the importance of PA for many dimensions of health, and its’ reported heritability, we sought to identify genetic variants that are associated with engagement in habitual physical activity, while considering important covariates such as season, physical activity at work, socio-economic status, and BMI.

In the UK Biobank, with a very large sample size and multiple measures of PA, we identified over 25 genome-wide significant SNPs. Although most were novel, some had previous links to disease or other traits. Among these, the variant in *CADM2*, a gene which encodes cell adhesion molecule 2, and is primarily expressed in the brain, has recently been linked to risk-taking behavior and other personality traits ^43–45^, as well as with information processing speed ^46^. The A allele at rs1376935 that we found to be associated with higher SSOE, is in LD in the European population with the G allele at the rs13084531 SNP which was found to be associated with risk-taking behavior ^45^. It thus appears that this locus may be important for several personality, cognitive, and behavioral traits, and may potentially be involved in reward systems. Additionally, it is important to note that this locus was significant for SSOE but not for the other traits. It may thus be specifically implicated in the proclivity to engage in intentional high-intensity exercise and sport, as opposed to more general and/or lower intensity PA.

Interestingly, a well-established variant in *APOE* (part of *APOE* ε4 allele), strongly implicated in Alzheimer’s disease ^47,48^, exhibited one of the strongest associations with PA, and remained significant upon meta-analysis. How the *APOE* risk allele is associated with greater PA is not clear. An exercise training study found that *APOE* ε4 carriers had a greater increase in aerobic capacity ^49^. This increased responsivity to PA could reinforce engagement in PA or be related to other factors that influence the tendency to engage in PA. Although another potential explanation for our finding is that individuals with a known family history of dementia or Alzheimer’s disease purposefully increase their levels of PA in the hope of reducing risk for developing the disease, our findings do not suggest that individuals with a first-degree family history of Alzheimer’s disease or dementia engage in higher levels of PA. It is important to note that the association between *APOE* and PA may lead to spurious gene-environment interactions ^50^, and thus further work is needed to clarify this observed association. Additionally, although the association remained after adjustment for BMI and other covariates, it remains to be determined whether some residual confounding might exist. Finally, it is important to note that none of the identified loci have previously been found to be associated with PA or related traits ^16,17^.

Previous studies have shown that BMI-associated genetic variants are also associated with PA ^51,52^. Similarly, we found an overall shared genetic basis for PA (especially accelerometer-based measures) with several obesity-related traits, suggesting that genetic risk for obesity coincides with genetic propensity for lower PA. There is likely a complex set of genetic, environmental, and phenotypic factors that connect PA and obesity across the lifespan, that involve many pleiotropic genetic factors.

Although the direction of correlation is reversed when BMI is included as a covariate, caution is warranted in interpreting results of genetic associations in which heritable covariates are included in the association model ^53^. Another major potential source of bias is collider bias, which occurs when one controls for a variable (i.e. BMI) that is caused by both another covariate (i.e. gene) and the outcome variable in the model (i.e. PA) ^54,55^.

Our study is strengthened by the large sample size, the availability of both self-reported and objective accelerometer-based measures of PA, and the availability of a replication cohort from a different country. However, we note several limitations. Given the relatively small genetic effect sizes observed for these PA phenotypes, we were insufficiently powered to formally replicate associations in the much smaller sample size in ARIC. Additional replication studies are thus needed to more robustly identify PA-associated loci. Furthermore, the self-report measures of PA used in ARIC differed from the one used in the UK Biobank. Both self-reported and accelerometer-based measures of PA are subject to various biases. Since both the UK Biobank and ARIC cohorts are comprised of middle-to late-middle-aged adults, the extent to which these results generalize to other age groups is not known. Furthermore, our results may not generalize to other ethnic/racial groups.

In conclusion, our study revealed several important new findings. Effect sizes were generally very small, given the very large sample size, the common variants identified, and the modest p-values. We identified over 20 variants, most of which were novel, and thus need further study. We identified a variant in *CADM2*, a gene previously been found to be associated with personality traits. We also identified a well-established major risk variant for Alzheimer’s disease in *APOE*, which was associated with higher levels of PA, suggesting the need for follow up studies to help clarify the nature of this observed association and its implication for understanding gene-environment interactions related to PA. We found genetic correlation of PA with obesity ^56,57^, psychiatric ^58,59^, educational ^60^, chronotype ^61^, and other traits. Genetic correlations with obesity may indicate extensive pleiotropy involving genes associated with both PA and obesity. The identification of genetic factors that predispose to high or low levels of PA will lead to a better understanding of the biological mechanisms underlying these proclivities. It may also lead to the identification of individuals less likely to engage in and/or adhere to PA, and consequently to the development of tailored behavioral strategies. Finally, the integration of genetic characteristics with lifestyle and environmental information may point to how lifestyle/environmental factors interact with genetic factors to influence levels of PA.

## Acknowledgments

This research was conducted using the UK Biobank Resource under Application Number 15678. We thank the participants and organizers of the UK Biobank. We also thank the participants and organizers of the ARIC study. Data from ARIC was obtained from dbGaP through accession number phs000280.v2.p1. The authors would like to acknowledge support from the National Institute of Diabetes and Digestive and Kidney Diseases grant (K01DK095032), the National Institute on Aging (AG019610), the State of Arizona and Arizona Department of Health Services (ADHS), and the McKnight Brain Research Foundation. The funders had no role in study design, data collection and analysis, decision to publish, or preparation of the manuscript.

## Atherosclerosis Risk in Communities

The Atherosclerosis Risk in Communities Study is carried out as a collaborative study supported by National Heart, Lung, and Blood Institute contracts (HHSN268201100005C, HHSN268201100006C, HHSN268201100007C, HHSN268201100008C, HHSN268201100009C, HHSN268201100010C, HHSN268201100011C, and HHSN268201100012C). Funding for GENEVA was provided by National Human Genome Research Institute grant U01HG004402 (E. Boerwinkle). The authors thank the staff and participants of the ARIC study for their important contributions.

## LDHUB Acknowledgements

We gratefully acknowledge all the studies and databases that made GWAS summary data available: ADIPOGen (Adiponectin genetics consortium), C4D (Coronary Artery Disease Genetics Consortium), CARDIoGRAM (Coronary ARtery DIsease Genome wide Replication and Meta-analysis), CKDGen (Chronic Kidney Disease Genetics consortium), dbGAP (database of Genotypes and Phenotypes), DIAGRAM (DIAbetes Genetics Replication And Meta-analysis), ENIGMA (Enhancing Neuro Imaging Genetics through Meta Analysis), EAGLE (EArly Genetics & Lifecourse Epidemiology Eczema Consortium, excluding 23andMe), EGG (Early Growth Genetics Consortium), GABRIEL (A Multidisciplinary Study to Identify the Genetic and Environmental Causes of Asthma in the European Community), GCAN (Genetic Consortium for Anorexia Nervosa), GEFOS (GEnetic Factors for OSteoporosis Consortium), GIANT (Genetic Investigation of ANthropometric Traits), GIS (Genetics of Iron Status consortium), GLGC (Global Lipids Genetics Consortium), GPC (Genetics of Personality Consortium), GUGC (Global Urate and Gout consortium), HaemGen (haemotological and platelet traits genetics consortium), HRgene (Heart Rate consortium), IIBDGC (International Inflammatory Bowel Disease Genetics Consortium), ILCCO (International Lung Cancer Consortium), IMSGC (International Multiple Sclerosis Genetic Consortium), MAGIC (Meta-Analyses of Glucose and Insulin-related traits Consortium), MESA (Multi-Ethnic Study of Atherosclerosis), PGC (Psychiatric Genomics Consortium), Project MinE consortium, ReproGen (Reproductive Genetics Consortium), SSGAC (Social Science Genetics Association Consortium) and TAG (Tobacco and Genetics Consortium), TRICL (Transdisciplinary Research in Cancer of the Lung consortium), UK Biobank. We gratefully acknowledge the contributions of Alkes Price (the systemic lupus erythematosus GWAS and primary biliary cirrhosis GWAS) and Johannes Kettunen (lipids metabolites GWAS).

